# Pseudo perfusion of Chinese Hamster Ovary (CHO) cells as a reliable platform for data generation to model and guide continuous perfusion biomanufacturing

**DOI:** 10.1101/2025.11.27.691016

**Authors:** Nikola G. Malinov, Shivam Barodiya, Marianthi G. Ierapetritou, Eleftherios T. Papoutsakis

## Abstract

Chinese Hamster Ovary (CHO) cell monoclonal antibody (mAb) production in continuous perfusion has witnessed a renewed interest within the biopharmaceutical industry. Widespread implementation of perfusion biomanufacturing, however, remains hindered by long process development timelines and high costs. Use of predictive scale-down platforms to generate large informative metabolic datasets and guide process development decisions is critical to decreasing a molecule’s time to market. While scale-down platforms based on the pseudo perfusion concept have been previously reported, they have not been rigorously validated. They are often limited by oxygen transport or insufficient metabolic characterization, reducing their role to a preliminary screening tool. Here, we report the design and validation of a pseudo perfusion platform based on a phenotype-driven approach to ascertain that the process emulates continuous perfusion characteristics and is not oxygen limited. Beyond metabolic and cell size steady state, we show that our pseudo perfusion design enables cell cycle subpopulation and intracellular antibody expression steady state. We also demonstrate that pseudo perfusion robustly predicts amino acid demands in continuous perfusion bioreactors with exceptional linear correlation across a broad range of cell-specific perfusion rates (CSPRs). When coupling the pseudo perfusion platform developed here with a workflow for metabolic characterization, we significantly augment the dimensionality and reliability of data which can be generated at this scale to gain actionable insights towards perfusion process design, ultimately reducing process development timelines and the associated costs.

**Highlights:** Residual lactate is a key proxy for oxygen transport in scale down platform design

Novel flow cytometry workflow confirms cell cycle and intracellular steady state

Pseudo perfusion robustly predicts metabolic phenotypes in continuous perfusion

K-means clustering analysis of nutrient rates provides insight into media design

## 1. Introduction

The growth of the biopharmaceutical industry has in large part been driven by recombinant protein production in Chinese Hamster Ovary (CHO) cells. CHO cells are the preferred host for recombinant protein expression given their ease of transfection, robust growth in chemically defined media, and innate ability to induce human-like post translational modifications [1]. Much of the manufacturing knowledge gained since the approval of the first therapeutic produced in CHO cells, tissue plasminogen activator (tPA) in 1987, has been transferrable across modalities and production platforms [2]. Monoclonal antibodies (mAbs) have since become the predominant class of biopharmaceuticals on the market today given their high target binding affinity and specificity, in turn revolutionizing healthcare by providing treatments for diseases ranging from cancers to autoimmune disorders [2].

Most CHO-based therapeutics that have emerged on the market are produced in fed-batch processes. Perfusion culture, which enables continuous upstream production, has numerous benefits over fed-batch including increased volumetric productivities due to higher cell densities, smaller facility footprints given reduced bioreactor volumes, and shorter product residence times leading to improved product quality profiles [3]. While the perfusion technology has existed for several decades, with the first commercial perfusion processes approved in 1992, widespread adoption of continuous cell culture has been constrained by several factors including the increased number of process parameters and resulting complexity of process design, the robustness of legacy fed-batch platforms, and the challenges of a changing regulatory landscape [3, 4]. Consequently, perfusion implementation has largely been case-specific, often reserved for labile biotherapeutics, such as Factor VIII which requires an exceedingly short residence time [5].

Scale-down perfusion platforms, with working volumes of several mL, are crucial for generating perfusion data to develop process models and inform design decisions. The ambr250 system is the preferred industry workhorse for scale-down studies, but comes with significant capital and operating costs [6]. An alternative non-instrumented scale-down platform is based on the concept of pseudo perfusion, in which the nutrient renewal and harvest are performed as an instantaneous medium exchange. The concept has been deployed with various culture vessels (ambr15 vessels, shake flasks, spin tubes, deep-well plates, and microwell plates) as a screening tool to assess cell clone performance, investigate media formulations, and inform process parameters such as the perfusion and cell-specific perfusion rates (CSPR) [6–14]. However, most studies have shown the need for additional larger-scale continuous perfusion experiments for process refinement given the disparity of metabolic phenotypes observed between pseudo and continuous perfusion at the same operating conditions. Moreover, the lack of direct pH and dissolved oxygen (DO) monitoring and control can lead to oxygen-limited and/or pH-stressed cultures, thereby constraining pseudo perfusion to serve only as preliminary screening tool rather than a platform to collect data with predictive potential for real perfusion processes.

Here, we advance the concept of pseudo perfusion as a reliable platform for process development by first applying a phenotype-driven approach to inform the selection of an appropriate vessel type, working volume, shake speed, and medium-exchange frequency to ensure sufficient oxygen transport to the culture and support industrially relevant cell densities exceeding 90×10^6^ cells mL^-1^. We demonstrate that pseudo perfusion enables steady state on four levels of increasing resolution. Cell growth, nutrient, and metabolite profiles along with cell diameter data confirm metabolic and cell-size steady state. Additionally, a novel workflow combining simultaneous cell cycle analysis and intracellular mAb heavy and light chain immunofluorescence staining was developed to demonstrate cell cycle subpopulation and intracellular mAb expression steady states. Subsequent validation with bench-scale continuous perfusion bioreactor cultures showed that pseudo perfusion supports nearly identical metabolic phenotypes across a range of process conditions.

## 2. Materials and Methods

### 2.1.1 Cell line and expansion

A CHO-K1 derived cell line (Clone A11 from the Vaccine Production Program Laboratory at the U.S. National Institute of Health) expressing the VRC01 IgG1 monoclonal antibody was used in this work. After the initial thaw, cells were passaged twice for inoculum preparation. Cultures were initiated at a 20 mL working volume in 50 mL vented spin tubes (Cell Treat Bio-Reaction Tube, Product Code: 229476) in HyClone ActiPro basal media (Cytiva) with 0.6 mM supplemental L-glutamine (Gibco). Cultures were grown in a NuAire CO2 humidified incubator (NU-5841) at 37 °C, 85% relative humidity, 5.0% CO2, and 20% O2 on a shaker plate (Thermo Fisher Scientific, Model No. 88881103) at 250 rpm shake speed, 90° angle, and 19 mm orbital throw. Cells were centrifuged at 180 × g for 5 minutes and resuspended in fresh basal media prior to inoculation to minimize carryover of inhibitory metabolites.

### 2.1.2 Pseudo perfusion culture

Cultures were seeded at 0.4×10^6^ cells mL^-1^ at a starting volume of 9.4 mL in HyClone ActiPro basal media with 0.6 mM supplemental L-glutamine in 50 mL vented spin tubes. 1.1 mL of culture broth was removed every day from days zero through three such that the working volume was reduced to 5 mL by day three after sampling. 0.1 mL of culture broth was used for cell counts while the remaining 1 mL was centrifuged at 200 × g for 5 minutes, 0.22 μm sterile-filtered, and saved at -80 °C for off-line metabolite analysis.

On day three, pseudo perfusion was initiated at an effective perfusion rate of 1 vessel volume per day (vvd^-1^) by performing 100% of the medium exchange every 24 hours or 50% of the medium exchange every 12 hours. For the medium exchange, cultures were centrifuged at 180 × g for 5 minutes, after which the supernatant was removed, and the cell pellets resuspended in pre-warmed perfusion media. An in-house perfusion medium formulation was adapted from an existing fed-batch protocol consisting of HyClone ActiPro basal media, 3% HyClone Cell Boost 7a feed media (Cytiva), 0.3% (v/v) HyClone Cell Boost 7b feed media (Cytiva), and supplemental glucose (Sigma Aldrich) targeting a final glucose concentration of 9 g L^-1^. 0.1 mL of resuspended culture broth was removed for validation cell counts during steady state experiments following the cell bleed. Cell growth continued until a pre-determined VCD setpoint, after which a daily intermittent cell bleed was initiated to maintain steady state at the desired setpoint for a minimum of seven days. Prior work has demonstrated that steady state at the transcriptomic and proteomic levels is achieved after seven days in continuous perfusion culture [15]. As the VCD increased, cultures were centrifuged and resuspended prior to the first set of cell counts to recover adhered cells. To further minimize cell adherence on the vessel walls, the cultures were transferred to new spin tubes every two to three days.

### 2.1.3 Continuous perfusion culture

Continuous perfusion experiments were performed in a 1 L Eppendorf BioFlo® 320 bioreactor at a working volume below 1 L. For cell retention, a hollow fiber filter module (Repligen) was operated in tangential flow filtration (TFF) mode at a target wall shear rate of 1100 s^-1^ with a centrifugal pump (PuraLev® i30MU, Levitronix, Switzerland). Two modules were used depending on the experiment: either a modified polyether sulfone (mPES) module of 20 cm length, 520 cm^2^ surface area, and 0.65 μm pore size or a polyether sulfone (PES) module of 20 cm length, 1000 cm^2^ surface area, and 0.2 μm pore size.

The bioreactor inoculation conditions were identical to the pseudo perfusion experiments. Batch mode operation ensued until day three, after which perfusion was initiated at a rate of 1 vvd^-^

Temperature was controlled at 37 °C. pH was controlled at 7.00 ± 0.05 via CO2 addition through a microsparger. Addition of base to control the pH was not needed across the conditions investigated in this work. Dissolved oxygen (DO) was maintained at 40% of air saturation. The sparging rate was allowed to vary between 0.02 – 0.40 SLPM using three-gas control (air, CO2, and O2), and a constant overlay at 0.05 SLPM of 100% air was used to maintain a positive pressure in the headspace. Using a pitch blade impeller, the agitation rate was set to 90 rpm at inoculation and increased in increments of 20 – 40 rpm to meet cellular oxygen demand at the set DO level. 5% Antifoam C Emulsion solution (Sigma Aldrich) was added as needed to reduce foaming. Daily samples were collected for offline measurements. During steady state operation, sample collection took place before and after the daily intermittent cell bleed.

### 2.2 Analytics

#### 2.2.1 Offline measurements

Viable cell density and percent viability were quantified using the trypan blue method on a DeNovix fluorescence cell counter. The average cell diameter was also determined using the DeNovix fluorescence cell counter. Glucose, lactate, ammonium, and choline concentrations were measured on a YSI 2950 bioanalyzer. Amino acid concentrations were measured on an Agilent HPLC 1260 Infinity II system with a precolumn derivatization protocol from Agilent. Briefly, primary amino acids were derivatized with OPA and secondary amino acids were derivatized with FMOC, separated on an Agilent Poroshell HPH-C18 column, and detected via a fluorescence detector [16]. MAb concentrations were also quantified on the Agilent HPLC 1260 Infinity II system using protein A chromatography as described previously [16]. A UV detection wavelength of 214 nm was applied to detect the lower daily titer concentrations in perfusion culture relative to fed-batch.

#### 2.2.2 Flow cytometry

During steady state experiments, 5×10^6^ cells from pseudo perfusion cultures were saved from the cell bleed and immediately before the subsequent medium exchange. 50 μL of culture, irrespective of the cell density and immediately before the subsequent medium exchange, were sampled during transient experiments to limit dilution effects on the VCD. Cells were washed twice with cold PBS (Gibco), fixed in 1 mL of 70% (v/v) ethanol, and stored at 4 °C until analysis. Published protocols for univariate cell cycle analysis using propidium iodide (PI) DNA staining and intracellular mAb heavy and light chain immunofluorescence staining were combined into a single and novel workflow outlined in the Supplementary File [17–21]. Cell cycle fractions were quantified with the FlowJo software. A subtractive model (Watson) was used to fit the G1 and G2 population peaks and estimate the S phase population. The G1 and G2 population peaks were constrained by an equal coefficient of variation (CV) to minimize the root mean squared deviation (RMSD) of the model fit.

### 2.3 Calculations

Specific rates for growth, nutrient consumption, and metabolite and mAb production were calculated for the pseudo and continuous perfusion bioreactor platforms used in this work and are outlined in Appendix A. Specific rates were compared across pseudo and continuous cultures to validate pseudo perfusion as a reliable scale-down platform.

## 3. Results and Discussion

### 3.1 Lactate depletion and industrially relevant viable cell densities support the absence of oxygen transport limitations in pseudo perfusion culture

Any non-instrumented scale-down platform will lack online monitoring and control of culture pH and DO. To address oxygen transport limitations, screening studies were performed to assess an appropriate culture vessel, working volume, and shake speed. Ranges for these parameters were informed by a comprehensive literature review of previous pseudo perfusion work. Other parameters, such as the rocking angle, was set to 90° based on literature findings [8], and the shaking diameter was fixed to 19 mm based on the constraints of the shaker plate used in this work. Selection of the final parameters followed a phenotype-driven approach guided by three target culture characteristics that support the absence of oxygen transport limitations. The first characteristic is the maximum VCD for a fixed perfusion rate and medium with no cell bleed, and concomitantly the critical CSPR (CSPRcritical), to be subsequently validated in bioreactor cultures. Decline of culture viability after nutrient depletion, which implies that the culture is nutrient limited rather than oxygen limited, is the second characteristic and substantiates the CSPRcritical determination. The final characteristic is the lactate shift from production to consumption, coupled with glucose and lactate depletion at high cell densities. Under oxygen-limited conditions, glycolysis presumably becomes the sole ATP production pathway, as there is insufficient oxygen to sustain ATP generation via oxidative phosphorylation and the electron transport chain, leading to lactate accumulation as a consequence [22]. Lactate accumulation due to oxygen limitation differs from lactate accumulation arising from carbon overflow metabolism during glucose overfeeding in the presence of sufficient oxygen, known as the Warburg effect [23, 24].

Spin tubes were selected as the final culture vessel following preliminary experiments with orbitally shaken T-flasks (Supplementary File). Spin tube pseudo perfusion control cultures were grown at a perfusion rate of 1 vvd^-1^ without a cell bleed at working volumes of 20 mL, 10 mL, and 5 mL, 250 rpm shake speed, and 19 mm orbital diameter to identify the working volume that would provide sufficient oxygen transport. The shake speed setpoint was selected to be at the upper range of reported values in the literature [25, 26] while operating below the maximum speed for the shaker plate used in this work. The 20 mL and 10 mL cultures were oxygen limited as evidenced by the significant lactate accumulation in between each medium exchange (Figure 1). In reducing the working volume from 20 mL to 10 mL, the maximum VCD increased, yet the lactate concentration remained high and achieved inhibitory levels as reported for CHO cells [23]. Moreover, the 20 mL and 10 mL cultures promoted an oxygen-limited steady state as would appear from the VCD profiles alone. Under these conditions, the cultures reached the oxygen saturation limit, which when coupled with sufficient nutrient supply as shown by the glucose profiles, and inhibitory lactate levels, will sustain a constant cell density without a cell bleed as previously reported for oxygen-limited continuous perfusion cultures [27]. In perfusion, a cell bleed is required to maintain true steady state culture even in instances where the process has been designed to minimize the specific growth rate at the desired operating conditions [28, 29]. Reducing the working volume to 5 mL increased the maximum VCD, followed by a pronounced reduction arising from the simultaneous depletion of several nutrients (Figure 1.a). Furthermore, the residual glucose and lactate for the 5 mL condition were depleted in between medium exchanges at high cell densities, implying a sufficiently oxygenated culture. During shaking, the whole 5 mL of the culture volume in the spin tubes forms a film on the vessel walls, hence increasing the surface area-to-volume ratio and improving oxygen transport.

**Figure 1:**
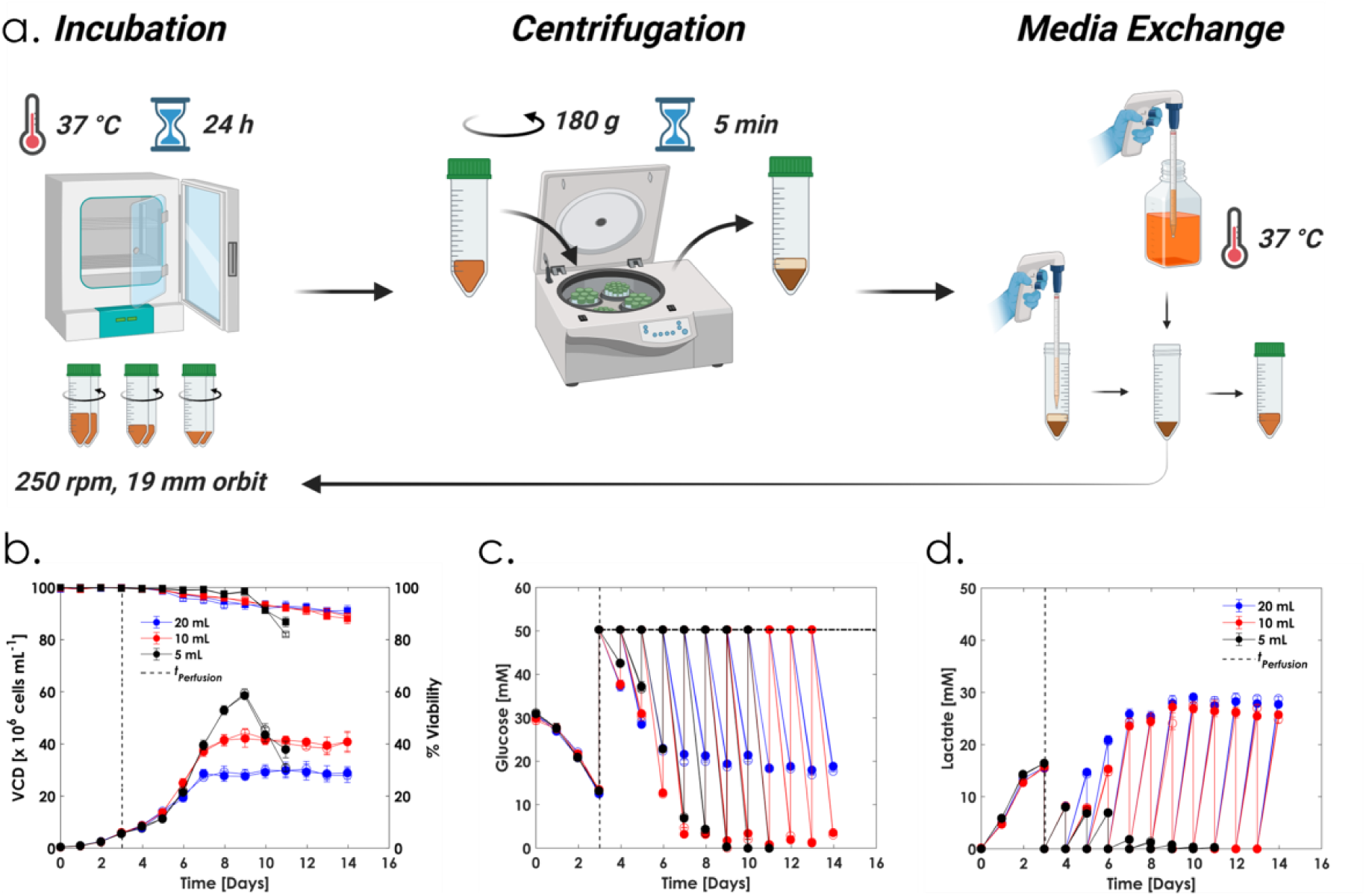
a) Pseudo perfusion culture at an effective perfusion rate of 1 vvd^-1^ with no cell bleed performed in spin tubes at 20 mL (blue), 10 mL (red), and 5 mL (black) working volumes, 250 rpm shake speed, and 19 mm orbital throw. **b)** VCD (circles) and percent viability (squares), **c)** residual glucose concentrations, and **d)** residual lactate concentrations. n=2 per condition with both biological replicates shown. Created with BioRender.com.

Perfusion processes routinely operate at cell densities nearing or exceeding 100×10^6^ cells mL^-1^ [30]. To push the limit of pseudo perfusion, we sought to demonstrate culture performance at similar cell densities by increasing the perfusion rate with the same medium formulation. Cultures were grown under dynamic perfusion conditions with no cell bleed following a ramping schedule for the perfusion rate increasing from 1, to 1.5, to 2 vvd^-1^ (Supplementary File, Figure S3.a). The maximum VCD for an effective perfusion rate of 2 vvd^-1^ with the medium in this study exceeded 90×10^6^ cells mL^-1^, with the residual glucose and lactate profiles demonstrating the desired characteristics indicative of sufficient oxygen transport (Supplementary File, Figure S3.b and S3.c). While the residual lactate concentration is nonzero, it is well below inhibitory levels and is completely depleted for one of the biological replicates by the end of the culture duration. Lactate accumulation occurred due to the dynamic feeding pattern, as the glucose concentration is reset to the medium concentration after every exchange, leading to partial overflow metabolism.

### 3.2 Pseudo perfusion enables metabolic, cell size, cell cycle subpopulation, and intracellular mAb expression steady state

It is critical that a scale-down perfusion platform can enable steady state culture, as in continuous perfusion bioreactors, to generate valuable data for metabolic characterization. Here, we demonstrate that the pseudo perfusion platform enables steady state on four levels of increasing resolution across a range of CSPRs. Metabolic steady state is observed by the VCD, nutrient, and metabolite profiles. Cell size steady state is reported by the average cell diameter and cell size distribution, indicating no drifts in cell physiology despite the intermittent operating conditions. We additionally developed a novel workflow combining simultaneous cell cycle analysis and intracellular mAb heavy and light chain immunofluorescence staining to confirm cell cycle subpopulation and intracellular mAb expression steady states.

Implementing an intermittent cell bleed will control the culture VCD about a desired setpoint and CSPR. We demonstrate pseudo perfusion’s versatility and reliability to investigate significantly different metabolic states by growing cultures at CSPRs of 50 and 25 pL cell^-1^ day^-1^ respectively (Figure 2.a and 2.b.). Given the low CSPR in the latter, an effective perfusion rate of 1 vvd^-1^ was maintained as a 50% medium exchange every 12 hours to approach continuous perfusion dynamics and allow stable CSPR control. The required bleed volume was calculated as described in Appendix A and applied to control the VCD setpoint and CSPR. As the cell bleed is discontinuous, a large bleed volume can induce a change in metabolism by shifting the culture to a new state. Figures 2.a-2.j demonstrate constant growth, nutrient consumption, and metabolite and mAb production profiles after each intermittent cell bleed across the steady state duration for each CSPR. Moreover, lactate is strictly produced, and ammonium is strictly consumed at each setpoint indicating that there is no change in metabolic state after each bleed (Figure 2.e-2.h). Furthermore, the residual concentration profiles for 19 of the 20 canonical amino acids support a strictly controlled metabolic steady state for each CSPR (Supplementary File, Figure S4). Therefore, despite the discontinuous medium exchange and cell bleed, pseudo perfusion can reliably generate steady state metabolic data.

**Figure 2:**
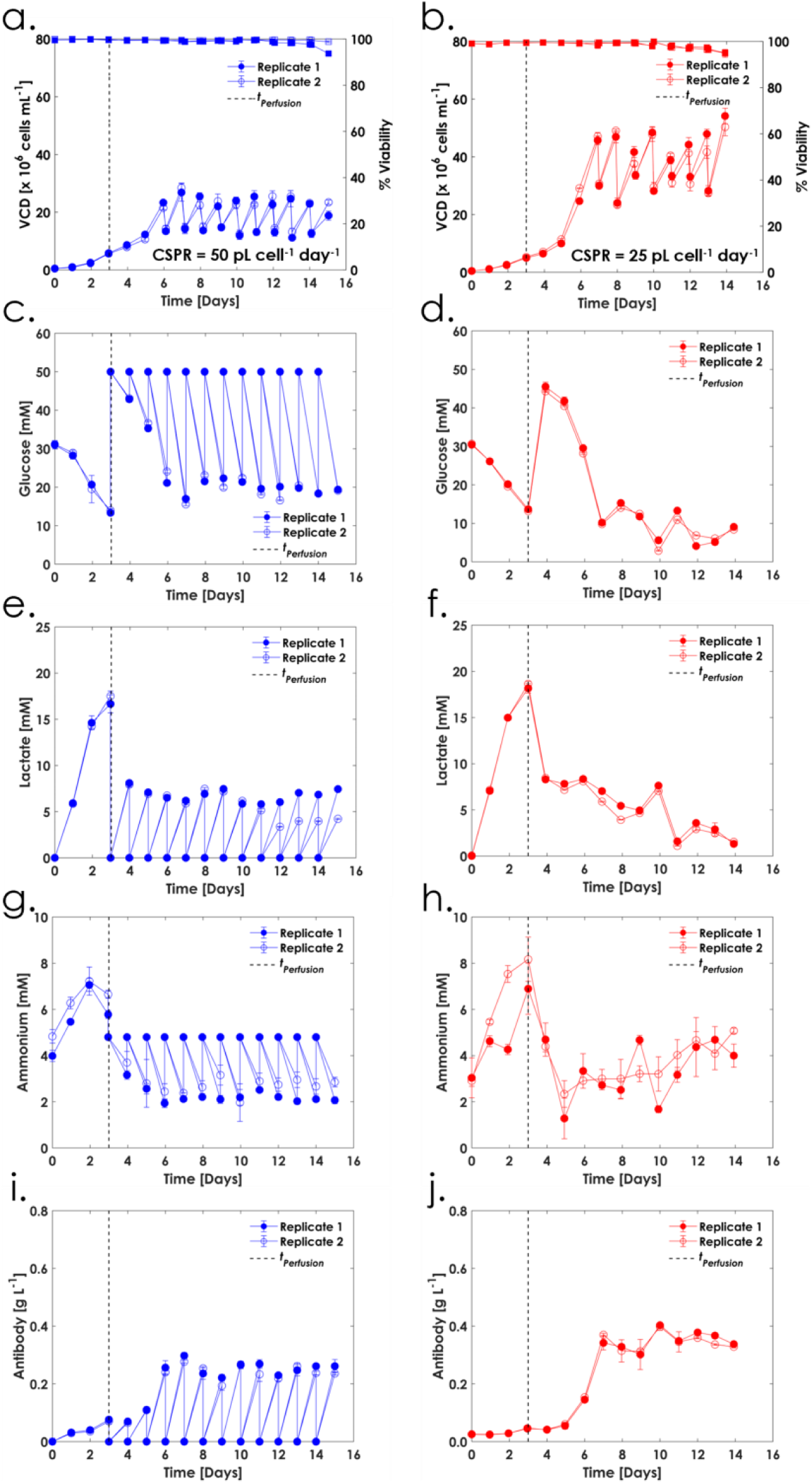
Engaging an intermittent cell bleed enables metabolic steady state in pseudo perfusion. VCD (circles) and % viability (squares) at CSPR setpoints of **a)** 50 and **b)** 25 pL cell^-1^ day^-1^ both with an effective perfusion rate of 1 vvd^-1^. **c)** and **d)** glucose, **e)** and **f)** lactate, **g)** and **h)** ammonium, and **i)** and **h)** antibody. n=2 per condition. For the CSPR at 25 pL cell^-1^ day^-1^, 1 vvd^-1^ was maintained as a 50% medium exchange every 12 hours with samples measured at only whole day intervals. Therefore, the residual concentrations at only whole day intervals are shown.

Steady state perfusion operation is largely based on VCD or CSPR setpoint control. Both will enable a steady state bioreactor environment, yet physiological changes can still occur, such as a drift in the average cell size [31], leading to viable cell volume setpoint control as a preferred alternative. Moreover, other studies have chosen to characterize and control their perfusion processes based on the biomass-specific perfusion rate (BSPR), defined as the volume of media consumed per volume of biomass [32]. The average cell diameter in pseudo perfusion was measured and shown to remain constant across the steady state duration for each CSPR (Figure 3.a, 3.c). There is a change in the cell size following each medium exchange and bleed due to the step change in culture conditions, however, the cell diameter range is constant over the steady state duration. Moreover, the relative cell size distribution as shown by forward scatter, remains constant at steady state (Figure 3.b, 3.d). Consequently, the dynamic operation of pseudo perfusion does not induce a drift in cell physiology which would otherwise not be detectable from measurements of the bulk culture properties alone such as VCD, glucose, and lactate concentrations.

**Figure 3:**
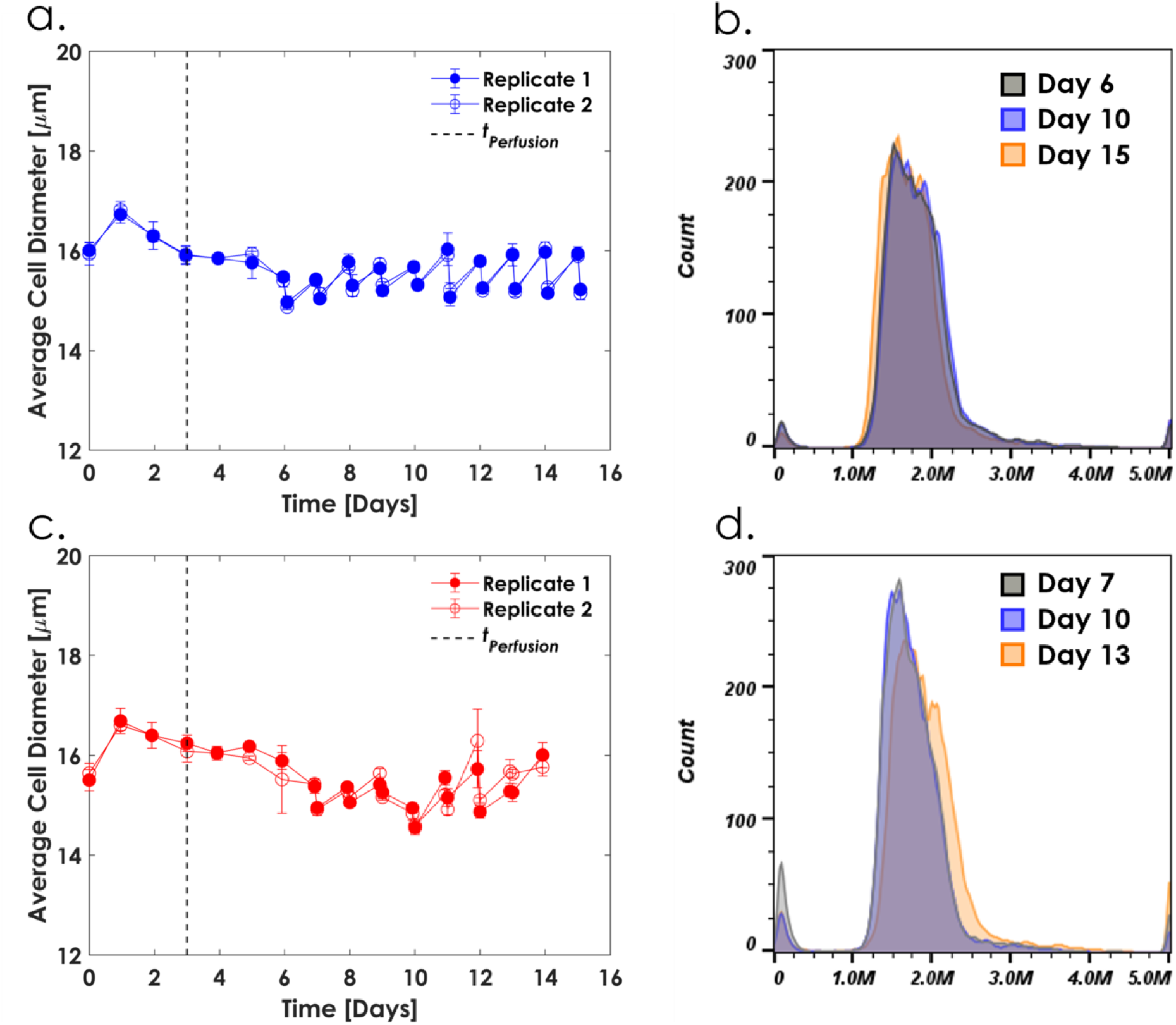
Average cell diameter and cell size distributions under steady state pseudo perfusion culture at CSPRs of **a)** and **b)** 50 and **c)** and **d)** 25 pL cell^-1^ day^-1^. n=2 per condition. The cell size distributions in **b)** and **d)** are reflective of ethanol-fixed cells. Therefore, the absolute cell size is not reflective of the true cell size at the time of sampling, however the relative distribution of cell sizes is preserved.

To further characterize the pseudo perfusion steady state, we combined univariate cell cycle analysis and intracellular mAb heavy and light chain immunofluorescence staining into a single novel workflow to gain insight into both subpopulation and intracellular dynamics. Univariate cell cycle analysis quantifies the fraction of cells in culture within each phase of the cell cycle; G1, S, and G2/M. Progression through the cell cycle is tightly regulated by cyclin-dependent kinases (Cdks) and cyclins (Cycs), with the complexation of phase-specific Cdks and Cycs enabling the cell cycle phase transitions [33]. At steady state and prior to each medium exchange, the distribution of cell cycle phases, or subpopulations, remains constant as also previously shown for continuous perfusion with a continuous cell bleed [28]. Figures 4.a and 4.b depict constant cell cycle fractions during the steady state duration for each CSPR, implying that despite the intermittent cell bleed and medium exchange, and the lack of pH and DO control, the distribution of cell cycle subpopulations remains unchanged. The larger variability in subpopulation fractions during steady state at the lower CSPR arises from momentary yet recoverable nutrient depletion in between medium exchanges (Supplementary File, Figure S4). When shifting to a lower CSPR, the majority subpopulation shifts from the S to the G1 phase. The G1 phase has been shown to have the longest duration of all cell cycle phases in CHO and other mammalian cells [34, 35], which is reflected by the lower specific growth rate of the culture (Figure 6.a). At a lower CSPR, achieved by operation at a higher VCD setpoint for a fixed perfusion rate, the nutrient availability per cell decreases promoting a decrease in the specific growth rate. Nutrient depletion in turn has been shown to promote cell cycle arrest in the G1 phase [36].

**Figure 4:**
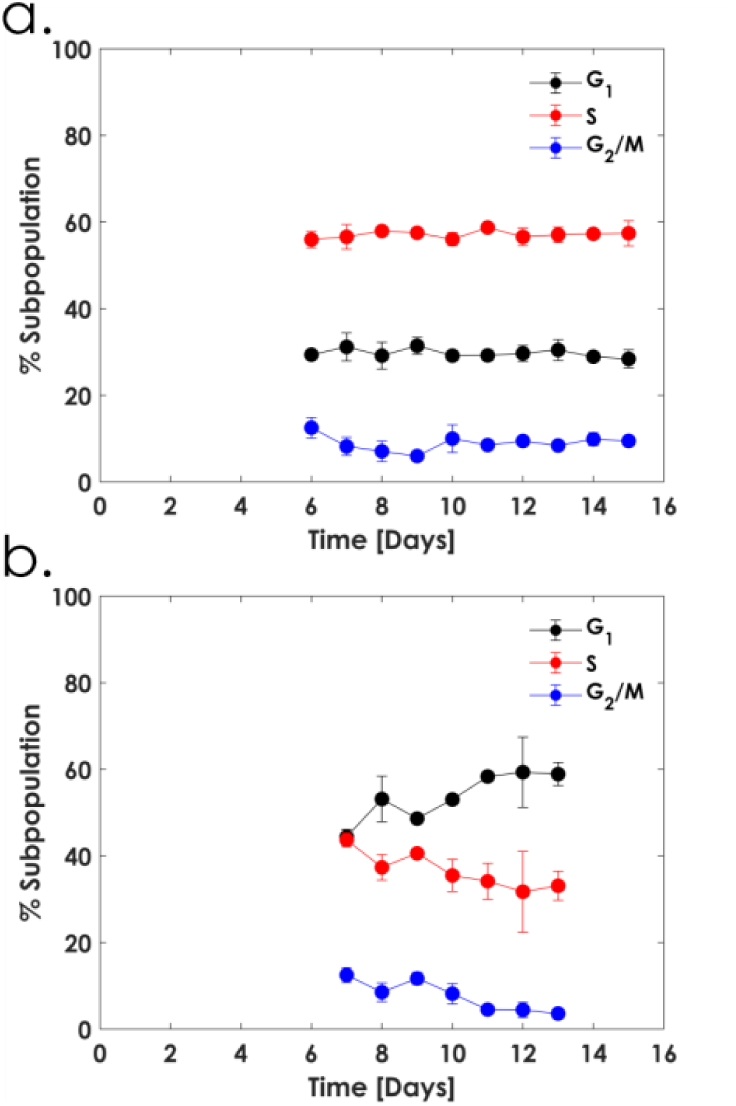
Cell cycle fractions from steady state pseudo perfusion at CSPRs of **a)** 50 and **b)** 25 pL cell^-1^ day^-1^. Values represent the average and one standard deviation of n=3.

Intracellular mAb heavy and light chain immunofluorescence staining is used during cell line development for the selection of high producing clones [20, 21]. Here, we used it as a method to interrogate steady state at the cellular level. Figure 5 depicts the emission peaks for the mAb heavy and light chains across the steady state duration for each CSPR to overlap with constant amplitude and spread. This is reflected by a constant geometric mean fluorescence intensity (MFI) during each day of steady state culture, implying intracellular steady state mAb production. Chain folding, assembly of the heavy and light chains into the mAb structure, and the initiation of the N-linked glycosylation process all occur in the rough endoplasmic reticulum (ER) [37]. The immunofluorescence staining protocol we used is specific to each of the mAb heavy and light polypeptide chains alone, which is as valid of a metric of cellular-level steady state as using immunofluorescence from antibodies targeting the heavy and light chains of a fully assembled mAb.

**Figure 5:**
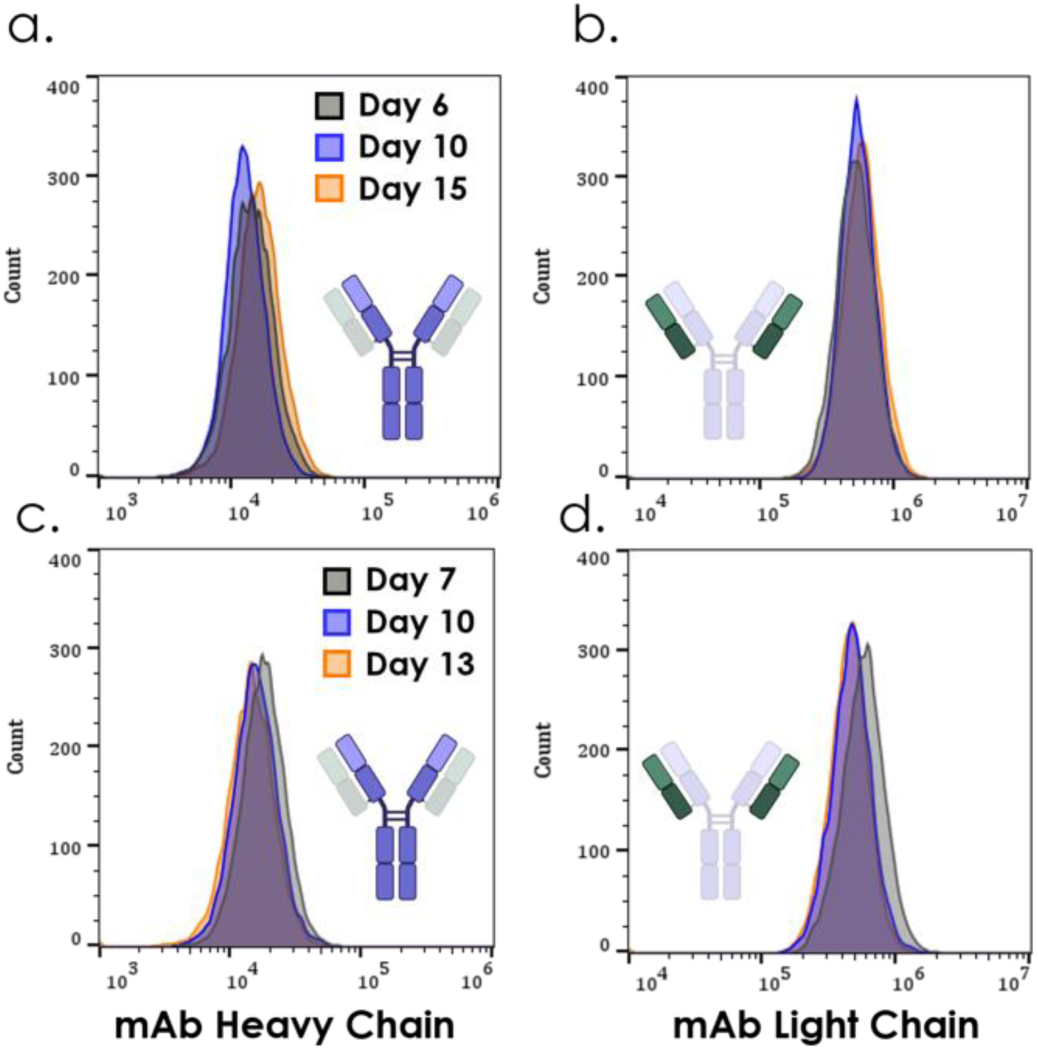
Intracellular mAb heavy and light chain immunofluorescence staining emission spectra from steady state pseudo perfusion at CSPRs of **a)** and **b)** 50 pL cell^-1^ day^-1^ and **c)** and **d)** 25 pL cell^-1^ day^-1^. Created with BioRender.com.

Taken together, the metabolic, cell size, cell cycle, and mAb-chain immunofluorescence data demonstrate that the dynamic operating conditions in pseudo perfusion do not prevent the cells from reaching and sustaining steady state. In our protocol, steady state culture is achieved because the intermittent operational time is kept below the characteristic timescale for each level of resolution: the doubling time at the metabolic and cell size level; the cell cycle phase-specific duration at the subpopulation level; and the post-translational residence time of the mAb chains in the ER at the intracellular level.

### 3.3 Pseudo perfusion predicts growth rates, metabolite and mAb production rates, and nutritional demands in continuous perfusion bioreactor cultures

After confirming that pseudo perfusion can promote steady state culture, we sought to validate that the resulting phenotypes are reflective of those observed in true continuous perfusion bioreactor cultures. Scale-down pseudo perfusion and continuous perfusion bioreactor cultures were grown at two steady states with target CSPRs of 50 pL cell^-1^ day^-1^ (VCD setpoint of 20×10^6^ cells mL^-1^) and 25 pL cell^-1^ day^-1^ (VCD setpoint of 40×10^6^ cells mL^-1^) respectively, and one dynamic condition with no cell bleed to reach the CSPRcritical, using the same medium formulation and fixed perfusion rate of 1 vvd^-1^. We performed a holistic comparative analysis of CHO-K1 VRC01 metabolic phenotypes between the 5 mL pseudo perfusion and 1 L continuous perfusion bioreactor process scales across CSPRs. We assessed the agreement between the specific growth rates, consumption of glucose and choline, production of lactate, ammonia, and mAb, and amino acid uptake rates.

Pseudo and continuous perfusion cultures exhibit similar specific growth rates across steady state and dynamic conditions (Figure 6.a). The agreement implies that the difference in cell retention modes, with centrifugation and resuspension in pseudo and TFF in continuous, do not confound the observed growth rates. As the cell density increases for a fixed perfusion rate, the nutrient availability per cell decreases leading to a reduction in the specific growth rate and in turn a decrease in all other specific rates. Moreover, comparison of the growth profiles between scales for the CSPRcritical case confirms that the pseudo perfusion platform in this work can sufficiently predict the CSPRcritical of 14-17 pL cell^-1^ day^-1^ for the medium formulation in this study (Supplementary File, Figure S5). Cell-specific productivities also agree across platforms within experimental conditions (Figure 6.b). While the cell-specific productivity appears to be growth-dependent, its reduction at lower growth rates is attributable to a decrease in the cell size (Supplementary File, Figure S8) confirming that growth and mAb production are not coupled.

The starkest difference between pseudo and continuous perfusion phenotypes lies in the specific lactate production rates at the two steady states despite the comparable glucose demands (Figure 6.c and 6.d). The larger specific lactate production rates in steady state pseudo perfusion arise from the dynamic feeding pattern. The glucose concentration is reset to the medium concentration after each medium exchange, promoting overflow of carbon metabolism [23, 24]. Under the CSPRcritical case at high cell densities, initial lactate production and subsequent consumption is observed prior to the following medium exchange (data not shown), hence the low residual lactate levels towards the end of the culture (Figure 1.d and S5.l). The low residual lactate concentration in high cell density continuous perfusion is promoted by the gradual addition of glucose, and thereby relatively low residual concentration in the bioreactor, which prevents overflow metabolism (Supplementary File, Figure S5.h and S5.i). Comparison of the VCD, viability, glucose, and lactate profiles across pseudo and continuous perfusion with no cell bleed demonstrates not only a comparable CSPRcritical of 14-17 pL cell^-1^ day^-1^, but nearly identical lactate phenotypes (Supplementary File, Figure. S5).

The medium in this study contains a low concentration of free ammonium which is consumed as a substrate under the two steady state conditions at similar rates between pseudo and continuous perfusion. Towards the end of the CSPRcritical condition, net ammonium production is driven by increased amino acid catabolism to support the higher cell densities in both pseudo and continuous perfusion, with productions rates in agreement across scales. Choline is a lipid precursor and therefore an important substrate for biomass proliferation and maintenance. The specific choline consumption rates agree between pseudo and continuous perfusion within experimental conditions (Figure 6.f), although choline is nearly or completely depleted in all cases by day six (Supplementary File, Figure S6).

Across all conditions, essential and nonessential amino acid specific consumption rates demonstrate statistically significant agreement between pseudo and continuous perfusion (Figure 7.a, 7.b, and 7.c). Glycine, which is observed to be strictly produced under the experimental conditions explored here, also demonstrates equivalent rates across the two process scales (Supplementary File, Figure S7). Upon lowering the CSPR, the cells exhibit a decreased amino acid demand, in alignment with the decreasing specific growth rate and concomitant increasing G1 phase majority subpopulation (Figure 4 and Figure S8.a). Beyond the robust predictions of amino acid requirements, the relative demand of each individual amino acid in comparison to all measured amino acids is preserved across CSPRs as seen by the relative location of each rate in Figures 7.a-7.c. To highlight this preserved relative demand, k-means clustering analysis was applied to each experimental condition to organize the consumption rates into similar groups. Across conditions, the rates constituting each cluster are largely conserved with only several rates shifting between clusters. The preserved grouping is in part attributed to the amino acid availability in the perfusion medium. This is apparent under the critical and 25 pL cell^-1^ day^-1^ CSPRs at which several amino acids are completely depleted, therefore presenting their maximum overall utilization rates. Interestingly, under nonlimiting conditions at a CSPR of 50 pL cell^-1^ day^-1^, the relative relationship between specific rates is also maintained. The cell cycle data in section 3.2 and Supplementary File clearly exhibit a significant change in metabolic state across experimental conditions, as also seen by the distribution of specific growth rates alone (Figures 6.a). Therefore, while the relative relationship between amino acid utilization rates is macroscopically conserved, the resulting intracellular metabolic flux distribution at each CSPR is undoubtedly different.

**Figure 6:**
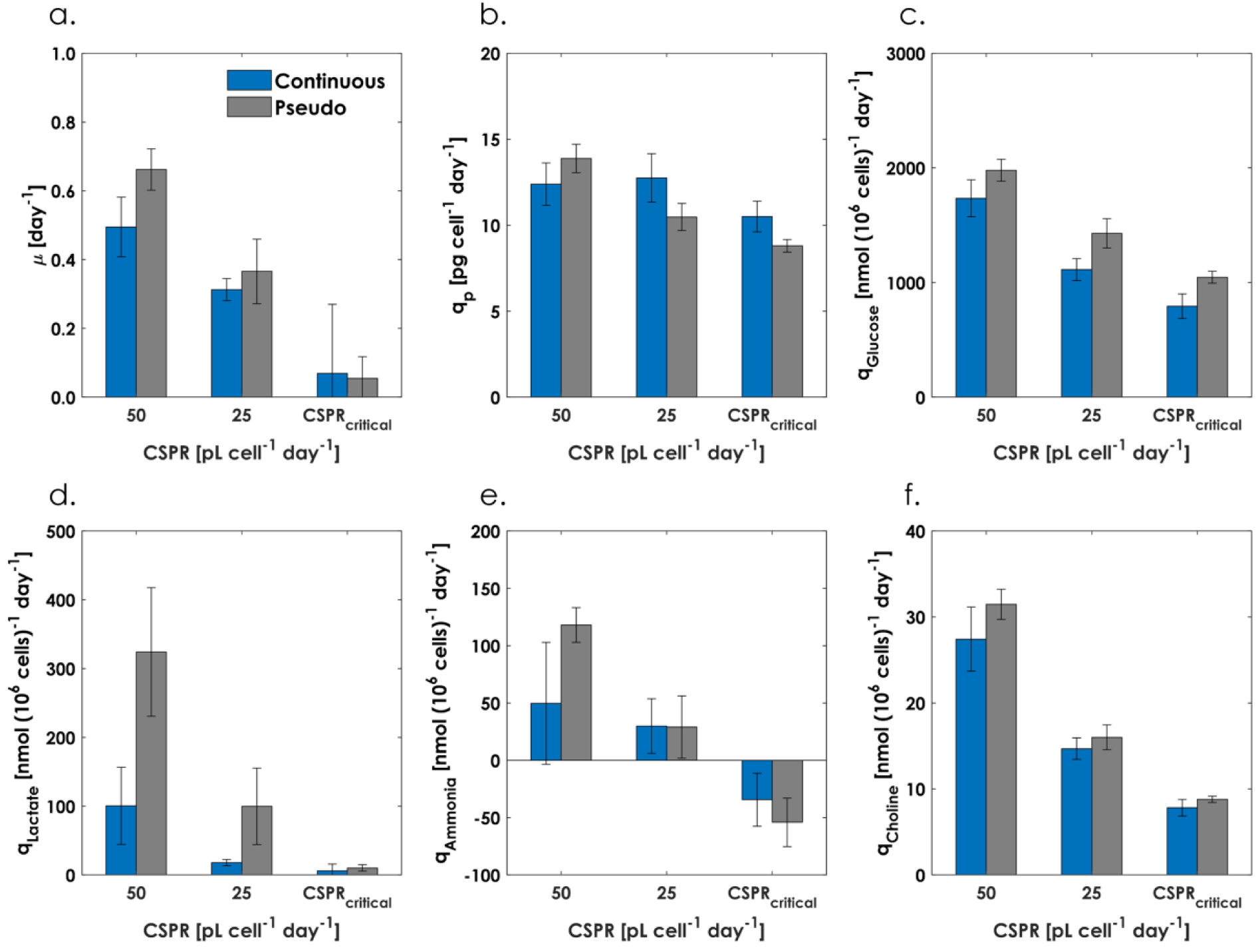
Comparison of the **a)** specific growth rates, **b)** cell-specific productivities, **c)** specific glucose consumption rates, **d)** specific lactate production rates, **e)** specific ammonium rates, and **f)** specific choline consumption rates between pseudo and continuous perfusion. For the CSPR of 50 pL cell^-1^ day^-1^, values represent the average and one standard deviation of the daily rates over the steady state duration. Steady state was maintained for nine days in pseudo and continuous perfusion. For the CSPR of 25 pL cell^-1^ day^-1^, values represent the average and one standard deviation of the daily rates over the steady state duration. Steady state was maintained for seven days in pseudo and three days in continuous perfusion due to filter fouling in the latter. For the CSPRcritical, values represent the average and one standard deviation of the daily rates over the two days during which the specific growth rate approaches zero near the maximum VCD. n=2 per scale and condition.

**Figure 7:**
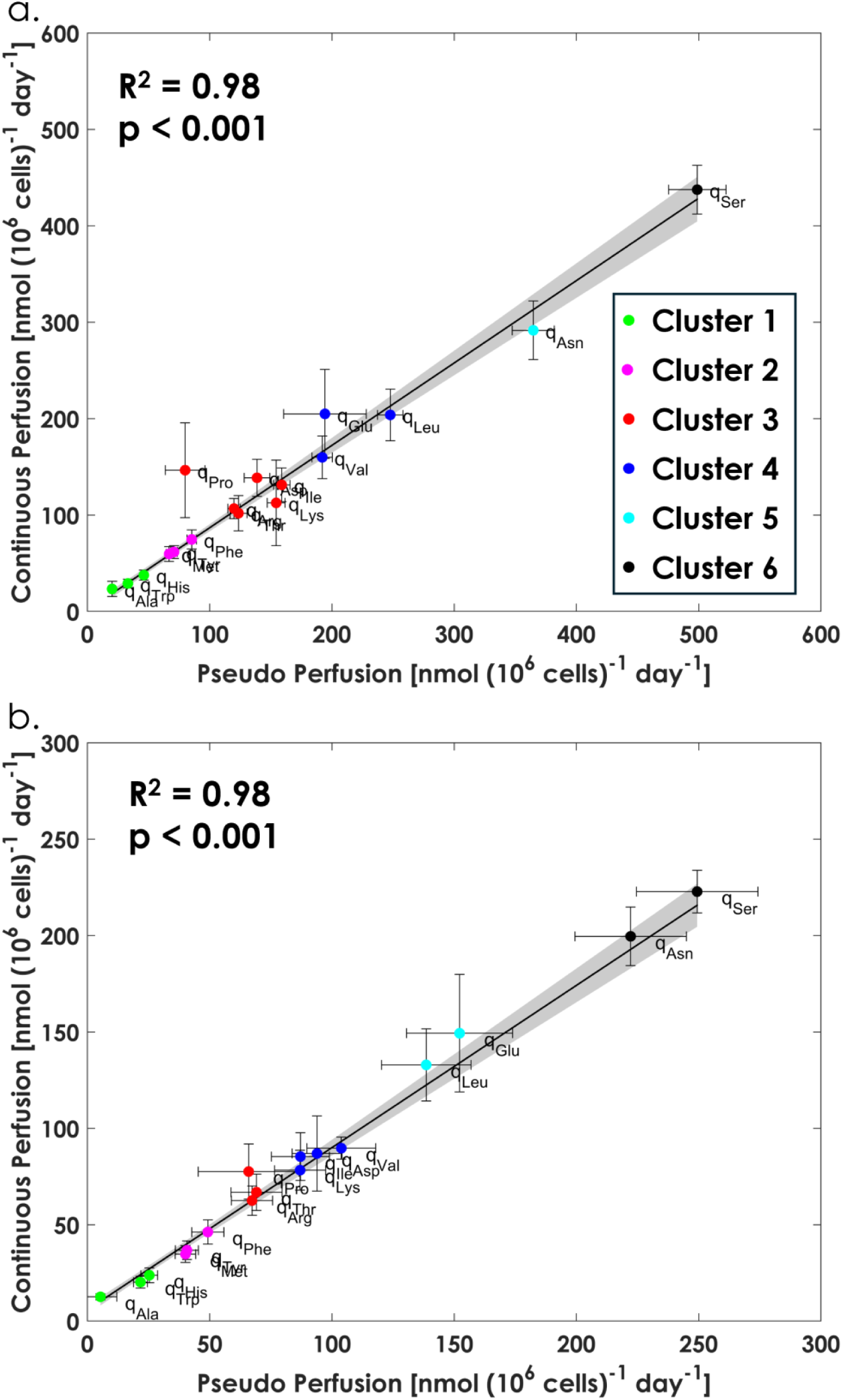
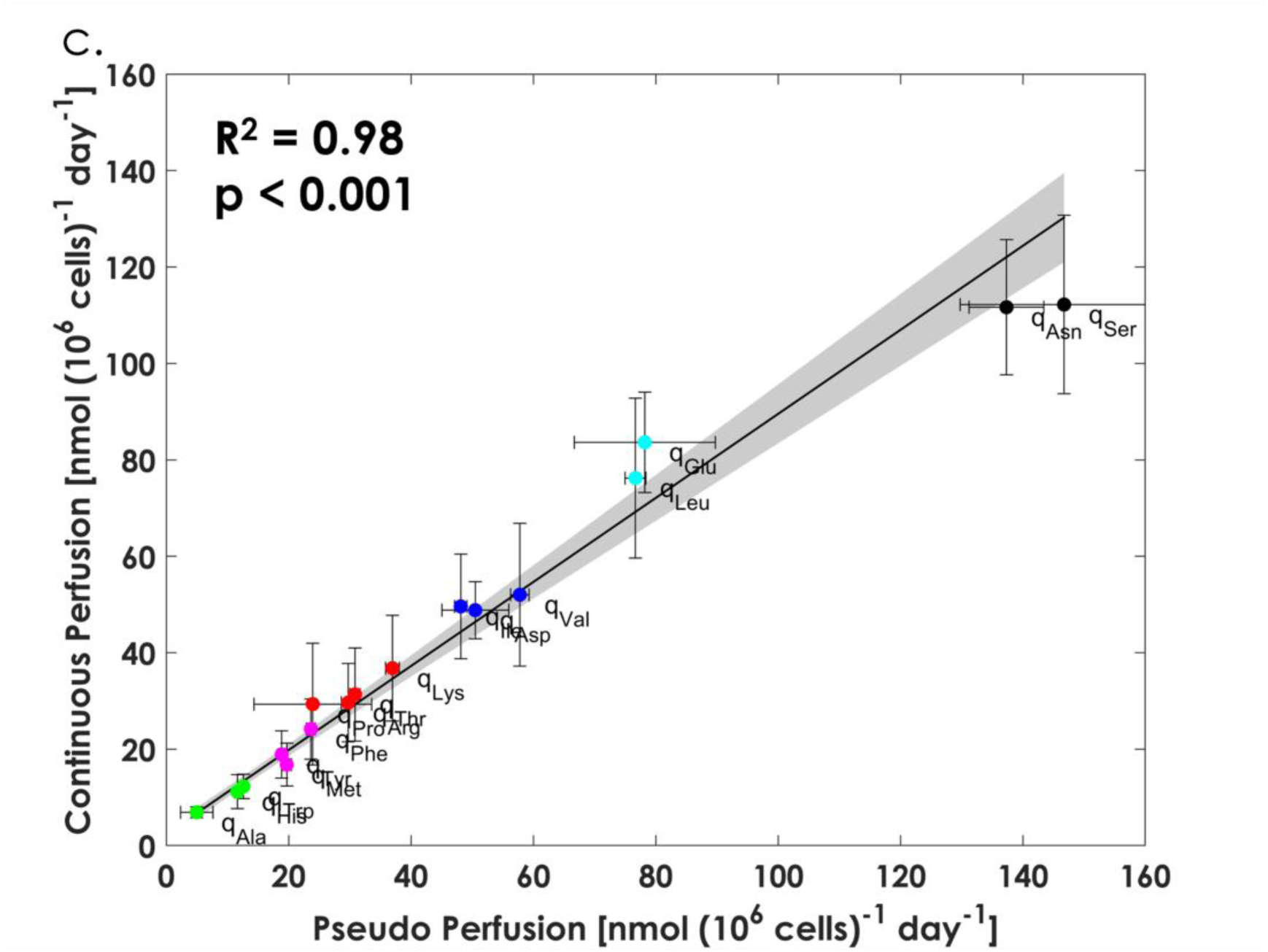
Specific amino acid consumption rates at: **a)** a CSPR of 50 pL cell^-1^ day^-1^, **b)** a CSPR of 25 pL cell^-1^ day^-1^, and **c)** at the CSPRcritical. Values and error bars are calculated as described for Figure 6. n=2 per scale and condition. The grey shaded area represents the 95% confidence interval for each linear model fit. For the k-means clustering analysis, the Elbow Method was applied first to select an initial range of k values. The Calinski-Harabasz index and Silhouette analysis were subsequently applied to determine an appropriate final k value.

In the context of process development, k-means clustering analysis of the pseudo perfusion specific rates presents a new application to points towards candidates that should be appropriately balanced in the medium or considered for concentrated feed strategies at the bioreactor scale. The amino acids with the largest consumption rates are also depleted at the CSPRcritical (data not shown), suggesting that their concentrations should be rebalanced to further reduce the CSPRcritical, or used as levers for steady state culture based on nutrient control. The k-means clustering analysis also reveals several interesting pairings of equivalent or near-equivalent amino acid uptake rates across all three conditions: tryptophan-histidine, methionine-tyrosine, threonine-arginine, isoleucine-aspartate, leucine-glutamate, and asparagine-serine, suggesting that their respective medium concentrations should be tightly controlled to satisfy the metabolic requirements of the cells while avoiding overfeeding and underutilization. Additionally, several of these pairings include poorly soluble amino acids, namely isoleucine and leucine. Therefore, analysis of pseudo perfusion specific rates can identify components with low solubility and high demand to preemptively mitigate challenges with large scale media preparation and storage required for bioreactor operation.

We show that pseudo perfusion data alone can accurately predict the CSPRcritical for a given medium, robustly predict the nutritional demands for continuous perfusion, and offer insight pertinent to media design and optimization that can be addressed with high fidelity at the small scale in subsequent rounds of pseudo perfusion experiments. The platform developed here can guide selection of media compositions, CSPRs, and bleed rates with minimal additional refinement at the continuous perfusion bioreactor scale. As the scope of this study is to demonstrate the reliability of pseudo perfusion as a small-scale predictive platform, inquiry as to the intracellular metabolism to explain the preserved rate clustering relationships and physiological relevance of the metabolic data is not pursued here. Application of constraint-based stoichiometric modeling, such as FBA and its variants, to investigate changes in intracellular metabolism in perfusion, offer a mechanistic explanation as to the preserved clustering, and suggest improved media formulations is a direction of future work.

## 4. Conclusions

Whether transitioning from traditional fed-batch to perfusion for an existing molecule or developing a new perfusion process for a novel product, application of reliable and predictive scale-down platforms that can generate high-quality and high-dimensional data are crucial in expediting process development timelines. In this work, we advance the pseudo perfusion concept by designing and validating a platform that can predict specific growth rates, nutrient demands, and metabolite and mAb production rates with a nearly perfect linear correlation to continuous perfusion bioreactor cultures across steady state and dynamic conditions. Apart from the agreement between scales, the high dimensional data generated from pseudo perfusion culture alone informs physiological and actionable insights towards continuous perfusion process design. Cell cycle analysis bolsters the importance and complexity of selecting an appropriate combination of media designs, CSPRs, and bleed rates to promote a desirable cellular state. The data and existing literature suggest further investigation into media components that sustain biomass maintenance and promote an increase in the average cell size at the selected operating conditions to boost cell-specific productivity. Additionally, the work discussed here speaks to the importance of incorporating higher resolution techniques, such as flow cytometric assays, in process characterization workflows. The information gain acquired from data on the individual cell level can provide insights into cellular state which are not detectable from standard bulk culture measurements.

The central limitations of any non-instrumented scale-down platform are the lack of pH and DO monitoring and control. Even with monitoring instrumentation coming to market, cost-effective pH and DO control at the pseudo perfusion scale remains a challenge. This work demonstrates the absence of oxygen transport limitations, yet the impact of uncontrolled pH is unknown. Offline measurements of the supernatant pH following each medium exchange under the CSPRcritical case spanned from 6.87 on day three to 7.81 by the end of the culture (data not shown). The lower end of the pH range can be attributed to the accumulated lactate at the conclusion of the batch regime, and the high end arises from the final ammonium accumulation. While this pH range does not significantly impact cell metabolism, as compared to continuous perfusion under the conditions investigated here, the effect on the mAb product quality has not been assessed. Studies with other cell lines and glycoproteins, however, have reported good agreement of glycosylation fractions and charge variants between pseudo and continuous perfusion even when the metabolic phenotypes showed disparity [6, 38].

Despite the above limitation, the work here substantially advances the metabolic information gain at the small scale and serves as a promising tool to expedite continuous perfusion process development timelines while also reducing the associated costs. The quality, dimensionality, and predictive reliability of the data that can be generated with this scale-down tool can also be coupled with metabolic modeling approaches to further aid design decisions. Constraint-based stoichiometric modeling can be applied to understand changes in intracellular metabolism and suggest improved media formulations. Pseudo perfusion data can also be leveraged to train kinetic or integrated metabolic models to simulate continuous perfusion operation, predict operating CSPRs under different media formulations, and reduce the number of larger scale continuous experiments needed for process optimization.

## CRediT authorship contribution statement

**N.G.M.:** Conceptualization, Methodology, Validation, Formal analysis, Investigation, Data curation, Writing – original draft, Writing – review & editing, Visualization. **S.B:** Investigation. **M.G.I.:** Conceptualization, Resources, Writing – review & editing, Supervision, Project administration, Funding acquisition. **E.T.P.:** Conceptualization, Resources, Data curation, Writing – review & editing, Supervision, Project administration, Funding acquisition.

## Appendix A

### A.1 Perfusion process parameters

The CSPR is defined according to Equation A.1:

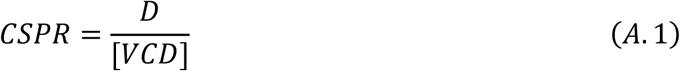

where [𝑉𝐶𝐷] is the viable cell density, 𝐷 is the perfusion rate defined as the volumetric flow rate of fresh perfusion medium introduced to the bioreactor, 𝐹, divided by the bioreactor working volume, 𝑉_𝑅_:

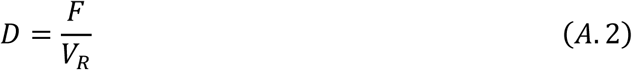

The critical CSPR, 𝐶𝑆𝑃𝑅_𝑐𝑟𝑖𝑡𝑖𝑐𝑎𝑙_, is the ratio of perfusion rate to viable cell density below which stable steady state operation is not possible.

### **A.2** Specific rates for pseudo perfusion

The time between discrete medium exchanges in the pseudo perfusion platform can be viewed as successive batch cultivations. Therefore, the specific rates, representative of the average rates over the period between medium exchanges, follow the form for batch culture. The specific growth rate, 𝜇, is calculated according to Equation A.3, assuming exponential growth:

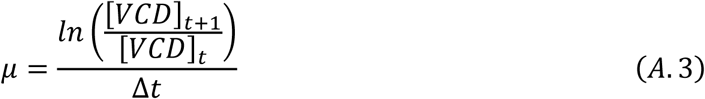

where [𝑉𝐶𝐷]_𝑡+1_ is the viable cell density at the time of the next medium exchange, [𝑉𝐶𝐷]_𝑡_ is the viable cell density at the time of the previous medium exchange, and Δ𝑡 is the time duration between complete medium exchanges which is maintained at one day intervals in this work. During steady state culture, an intermittent cell bleed is applied. The required daily bleed volume,

𝑉_𝑏𝑙𝑒𝑒𝑑_, to maintain the VCD setpoint, [𝑉𝐶𝐷]_𝑠𝑝_, is calculated from a mass balance on the viable cells:

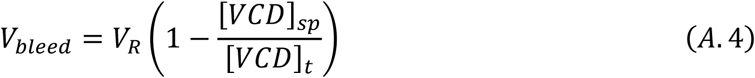

The specific rates for substrate consumption, 𝑞_𝑆_, which include glucose, choline, and amino acids, are also calculated from the batch culture mass balance applied in between medium exchanges. The substrate concentration after each medium exchange is assumed to be reset to that in the perfusion medium, [𝑆]_𝑚𝑒𝑑𝑖𝑎_:

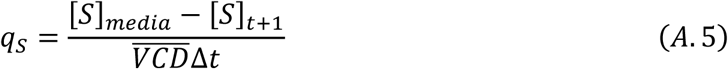

where 𝑉̅̅𝐶̅̅𝐷̅̅ is the average cell density in between medium exchanges, and [𝑆]_𝑡+1_ is the substrate concentration at the timepoint immediately before the next medium exchange.

The specific rates for metabolite and product generation, which include lactate, ammonium, and mAb are calculated according to Equation A.6 assuming that the metabolite and product concentrations after each medium exchange are reset to zero given their absence in the perfusion

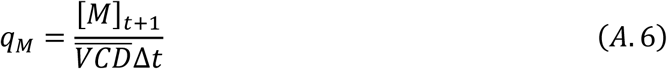

where [𝑀]_𝑡+1_ is the metabolite or product concentration at the timepoint immediately before the next medium exchange.

Existing batch and fed-batch protocols for the CHO-K1 VRC01 cell line supplement 0.6 mM glutamine upon inoculation, which is typically depleted by day three of culture at the end of the batch period [16, 39]. The pseudo and continuous perfusion culture protocols developed in this work follow the same inoculation and initial three-day batch regime. As such, by day three the glutamine is depleted (Supplementary File, Figure S4). While the glutamine degradation rate into pyrrolidone carboxylic acid and ammonium in ActiPro basal media has been experimentally determined [40], it is not included in the specific glutamine consumption rate or the specific ammonium rate calculations given the absence of glutamine in both the culture from day three onwards and the perfusion medium itself. All analysis in this work is performed after the initial batch regime.

For pseudo perfusion cultures in which the effective perfusion rate of 1 vvd^-1^ was performed as a 50% medium exchange every 12 hours, offline analysis for most nutrients and metabolites was completed on whole day intervals only. This was done given the large number of samples from both the increased sampling frequency and biological replicates. The time points at whole day intervals, however, are still representative of a 100% medium exchange, therefore Equations A.3, A.5, and A.6 were applied to calculate 𝜇, 𝑞_𝑆_, and 𝑞_𝑀_ respectively.

### **A.3** Specific rates for continuous perfusion

While the perfusion bioreactor is operated with an intermittent cell bleed, the perfusion medium addition and harvest removal do occur continuously. Therefore, 𝜇 in continuous perfusion is calculated according to Equation A.3 given the absence of a continuous cell bleed. The bleed volume required for the daily intermittent cell bleed at steady state is calculated from Equation A.4.

The specific substrate consumption rates are calculated from the continuous perfusion mass

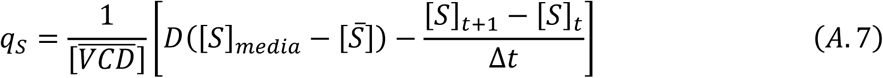

where [𝑆̅] is the average substrate concentration in the bioreactor in between sampling times maintained at one day intervals in this work.

The specific metabolite and product generation rates are calculated similarly:

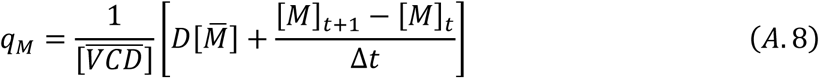

where [𝑀̅] is the average metabolite or product concentration in the bioreactor in between sampling times.

## Data Availability Statement

Data will be made available upon reasonable request.

