## Supplementary File for "Pseudo perfusion of Chinese Hamster Ovary (CHO) cells as a reliable platform for data generation to model and guide continuous perfusion biomanufacturing"

1. **Continuous perfusion culture**

Continuous perfusion at higher cell densities was faced with filter fouling in the TFF retention module as demonstrated by the product sieving profile (Figure S1). Product sieving, $PS$, is calculated according to Equation S1

$$\begin{aligned} PS=\frac{\left[ mAb \right]_{Permeate}}{\left[ mAb \right]_{Bioreactor}}\times100\%\#\left( S1 \right) \end{aligned}$$

where $\left[ mAb \right]_{Permeate}$ is the mAb concentration in the permeate and $\left[ mAb \right]_{Bioreactor}$ is the mAb concentration in the bioreactor.

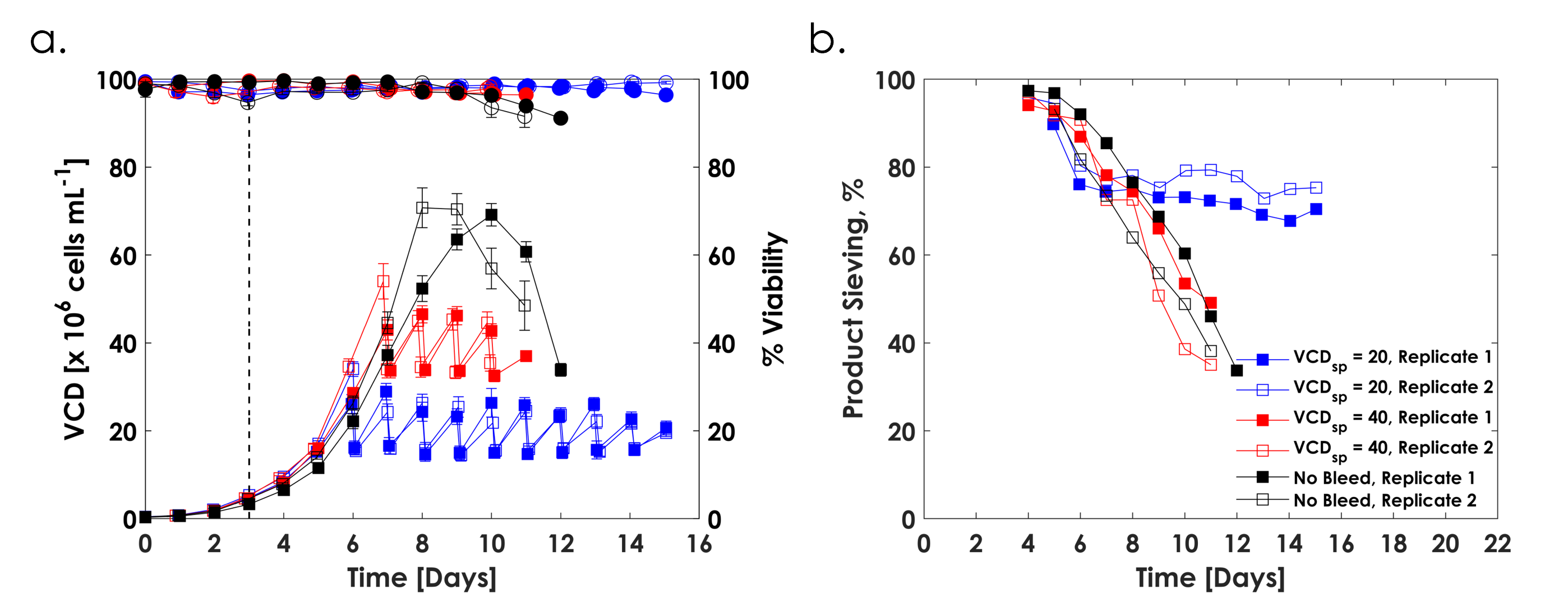

**Figure S1: a)** VCD (squares) and percent viability (circles) in continuous perfusion culture at a VCD_sp_ of 20×10^6^ cells mL^-1^ (blue), a VCD_sp_ of 40×10^6^ cells mL^-1^ (red), and No Bleed (black). Values and error bars represent the average and one standard deviation of ten technical replicates. **b)** Corresponding product sieving profiles. n = 2 per condition with both biological replicates shown.

Prior to filter fouling, steady state at the VCD_sp_ of 40×10^6^ cells mL^-1^ was sustained for three days. Under the No Bleed condition, culture decline due to nutrient starvation outpaced the onset of filter fouling. Therefore, both cases are reflective of the innate cellular state imposed under their respective conditions prior to any disturbances arising from retention module fouling.

2. **Flow cytometry**

Fixed cells were washed twice with 200 μL of cold PBS at 1000 × g for five minutes. A third wash with 200 μL of 5% (v/v) Fetal Bovine Serum (FBS) (Gibco) in cold PBS was done to block potential cross-reactive antibodies. Cells were resuspended in 200 μL of mAb heavy and light chain staining solution containing 0.1% (v/v) Triton X-100 (Sigma-Aldrich) in PBS, 5% FBS, and fluorescently labeled secondary antibodies diluted 1:64. Goat anti-Human IgG Fc Cross-Adsorbed secondary Antibody, DyLight 755 (Thermo Fisher) was selected for the $\gamma$ heavy chain. Goat anti-Human IgG Recombinant secondary antibody, Alexa Fluor Plus 488 (Thermo Fisher) was selected for the $\kappa$ light chain. After a 1-hour incubation period at 37°C in the dark, cells were washed in 200 μL of PBS at 1000 × g for five minutes and resuspended in 1 mL of PI staining solution containing 10 μg mL^-1^ PI (Thermo Fisher), 100 μg mL^-1^ RNase A (Thermo Fisher Scientific), and 0.1% Triton X-100 in PBS. After a 10-minute incubation at 37°C in the dark, samples were transferred to flow cytometry tubes for analysis.

All samples were analyzed on a CytoFLEX S flow cytometer (Beckman Coulter). The debris field was gated on the side scatter (SSC) vs forward scatter (FSC) plot. Doublets were gated as the population laying above the diagonal of the 690/50 bandpass filter on the pulse area versus height plot, while singlets were gated as the population laying along the diagonal. 10,000 events were recorded in the singlets gate for every sample. Negative controls were previously run to establish the workflow and determine if compensation was necessary. The 690/50 bandpass filter was used to assess DNA staining by PI for cell cycle analysis. The 780/60 bandpass filter was used for heavy chain analysis, and the 525/40 bandpass filter was applied for light chain analysis.

3. **Preliminary screening studies**

First, orbitally shaken T75 flasks were investigated as a pseudo perfusion platform. The rationale is that the surface area-to-volume ratio is significantly increased, relative to other orbitally shaken vessels, improving culture oxygenation. The working volume was set to 10 mL based on previous work demonstrating the interdependent relationship between liquid level height, oxygen transport, and lactate metabolism for hematopoietic cells cultured in flat-bed bioreactors [1]. The pseudo perfusion cultures were grown at 150 rpm shake speed and 19 mm orbital diameter with no cell bleed. To facilitate the medium exchange, the entire culture was transferred to a 50 mL spin tube, processed as outlined in section 2.1.2, and transferred back to the T75 flask.

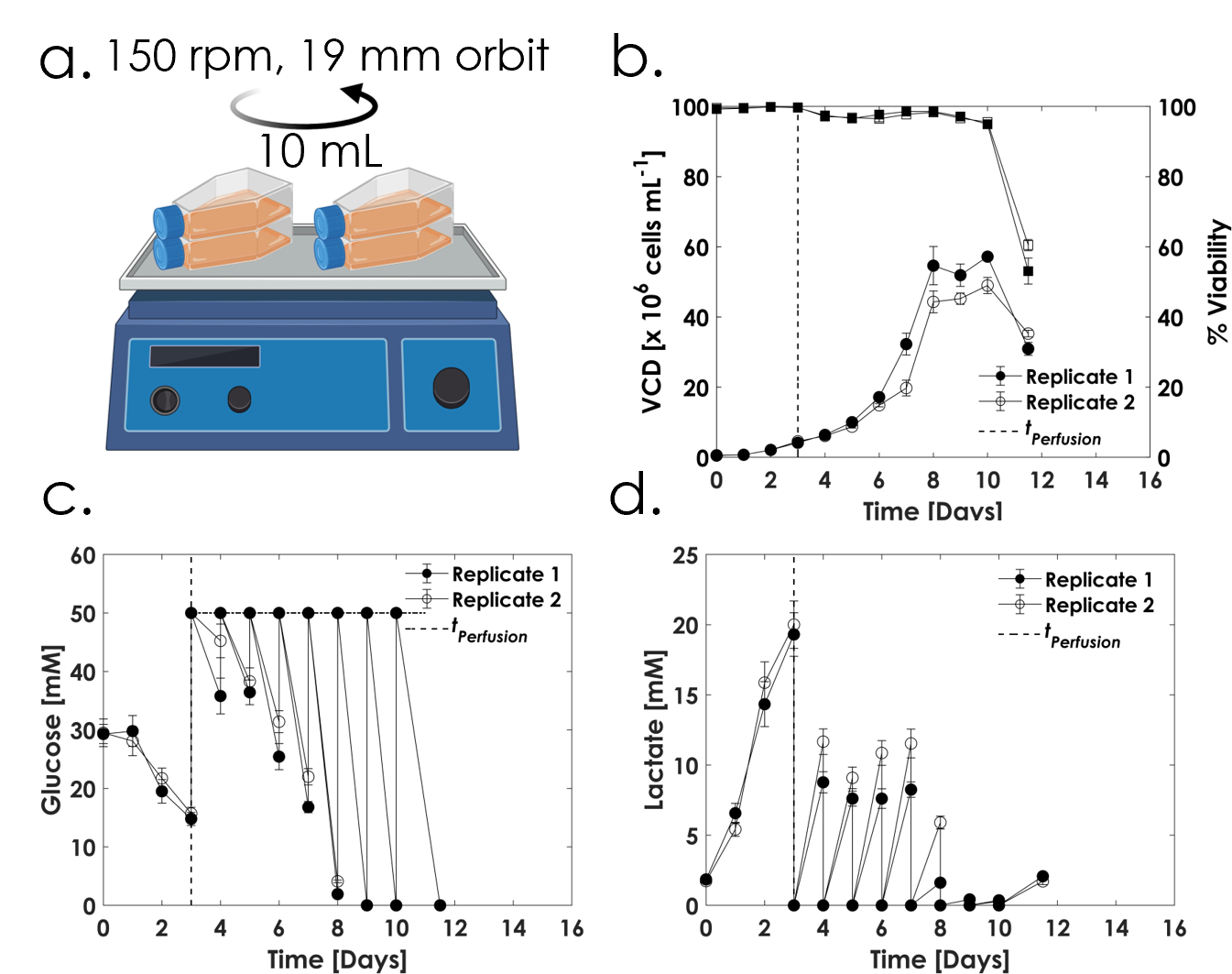

**Figure S2:** **a)** Pseudo perfusion culture, at an effective perfusion rate of 1 vvd^-1^ with no cell bleed performed in orbitally shaken T75 flasks at a 10 mL working volume, 150 rpm shake speed, and 19 mm orbital throw. **b)** VCD (circles) and percent viability (squares), **c)** residual glucose concentration, and **d)** residual lactate concentration. n=2 with both biological replicates shown.

Culture performance demonstrated all three characteristics which support the absence of oxygen transport limitations (Figure S2). The maximum VCD and minimum CSPR were within range of reported values achieved in continuous perfusion culture, offline metabolites analysis revealed depletion of glucose and several amino acids (data not shown), and the residual lactate profile showed the desired metabolic shift and eventual depletion. However, the vessel geometry induced a propensity for the accumulation of unrecoverable adhered cells on the vessel walls as the culture progressed, giving way to a misleading growth profile. With a fraction of the cells adhered to the vessel wall, yet sill covered by the culture broth during shaking, the measured VCD was an underestimate of the true VCD. As such, the T75 flasks were replaced with spin tubes moving forward. Nonetheless, the initial study demonstrated the proof-of-concept in applying the residual lactate concentration as a proxy for oxygen transport limitations.

4. **Dynamic pseudo perfusion enables industrially relevant cell densities**

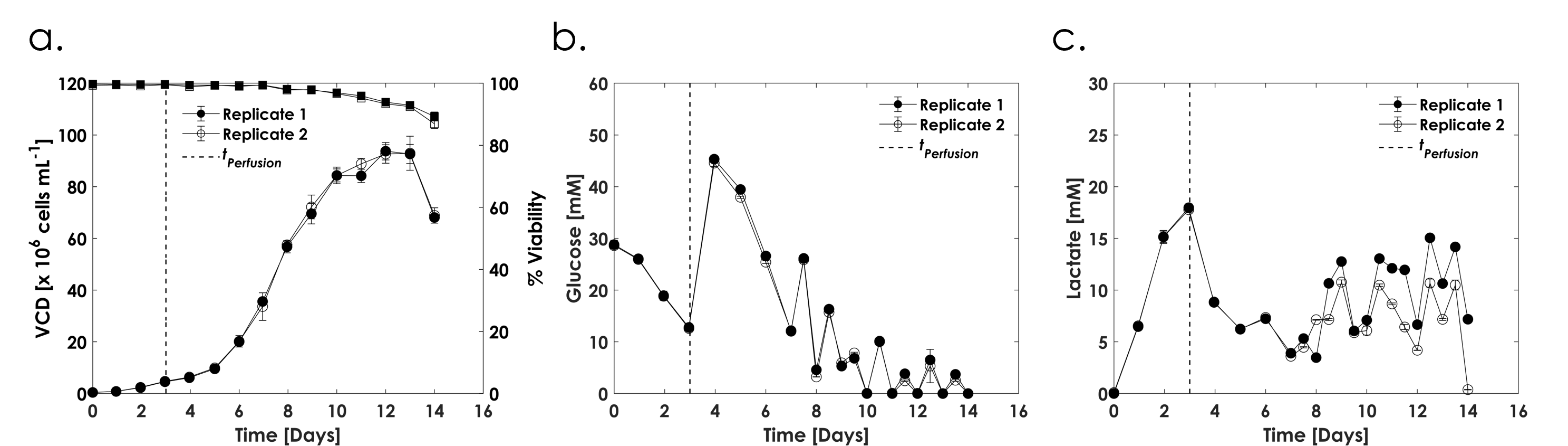

**Figure S3: a)** VCD (circles) and percent viability (squares), **b)** residual glucose concentration, and **c)** residual lactate concentration for dynamic pseudo perfusion with no cell bleed. Pseudo perfusion was initiated on day three at an effective perfusion rate of 1 vvd^-1^, 1.5 vvd^-1^ between days seven and eight, and 2 vvd^-1^ from day eight onwards. Concentrations only before each medium exchange are depicted in **b)** and **c)** for figure clarity. n =2.

5. **Metabolic steady state in pseudo perfusion**

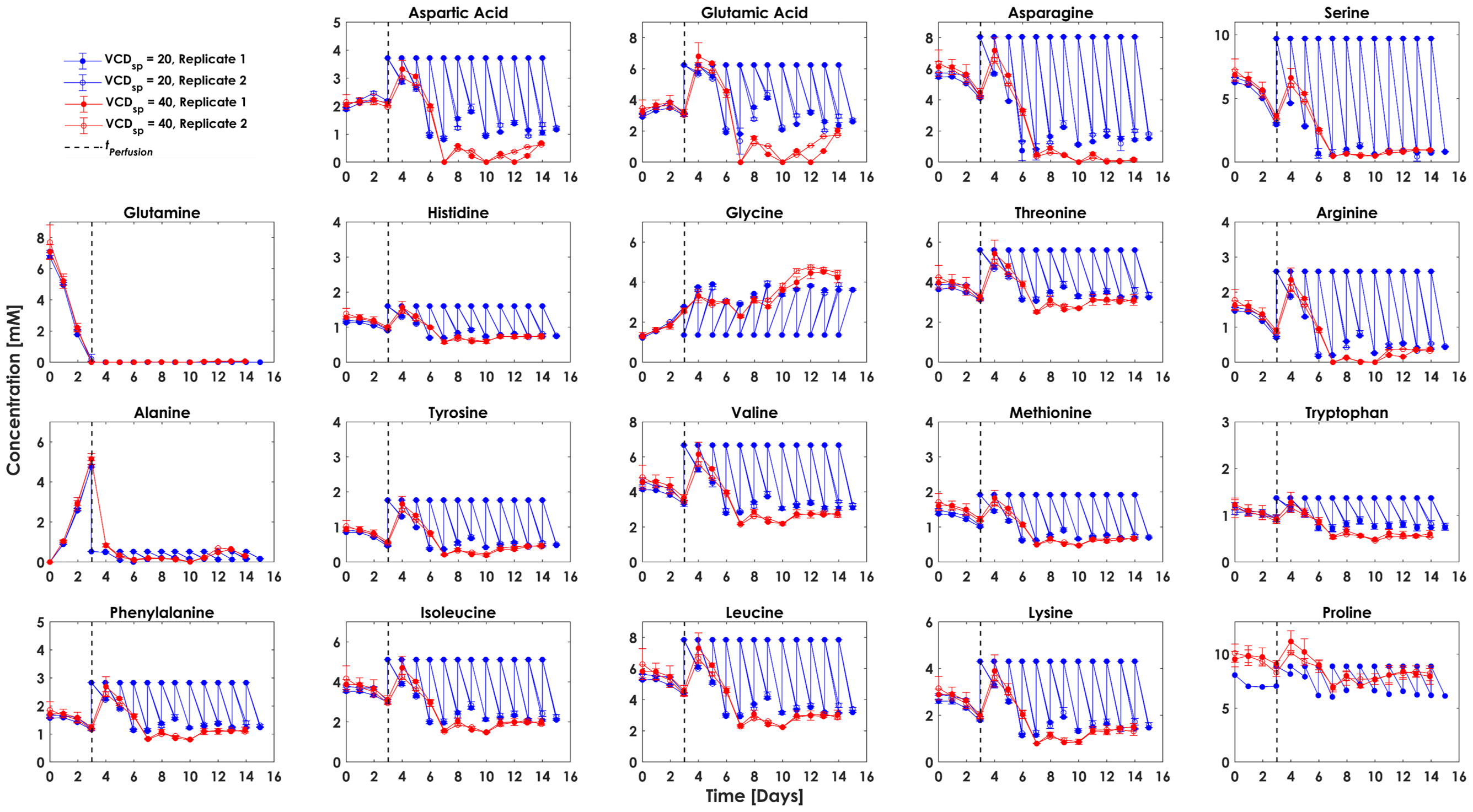

**Figure S4:** Amino acid concentrations under steady state pseudo perfusion at CSPRs of 50 (blue) and 25 pL cell^-1^ day^-1^ (red). For the latter, 1 vvd^-1^ was maintained as a 50% medium exchange every 12 hours with samples measured at only whole day intervals. Therefore, the residual concentrations at only whole day intervals are shown. n=2 per condition.

6. **Growth, glucose, and lactate profiles in pseudo and continuous perfusion**

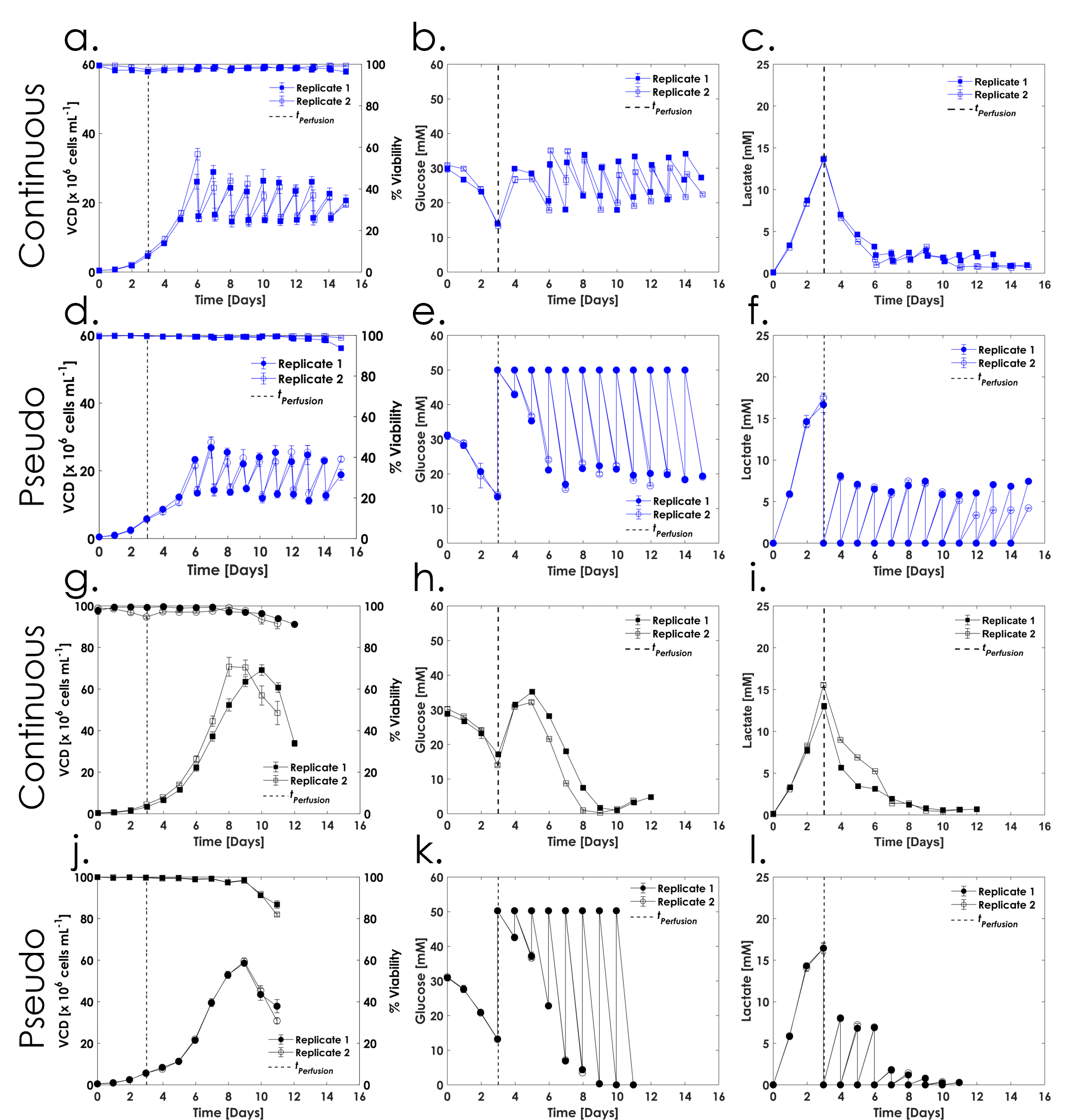

**Figure S5:** VCD, percent viability, residual glucose, and residual lactate concentration profiles for **a), b)** and **c)** continuous perfusion and **d), e)** and **f)** pseudo perfusion at a VCD_sp_ of 20×10^6^ cells mL^-1^. **g), h)** and **i)** continuous perfusion and **j), k),** and **l)** pseudo perfusion with no cell bleed. n=2 per scale and condition. The oscillations in the continuous profiles arise from the intermittent cell bleed. The oscillations in the pseudo perfusion profiles arise from the medium exchanges.

7. **Choline profiles in pseudo and continuous perfusion**

**Figure S6:** Residual choline concentrations in **a)** pseudo perfusion and **b)** continuous perfusion culture across all conditions. n = 2 per scale and condition.

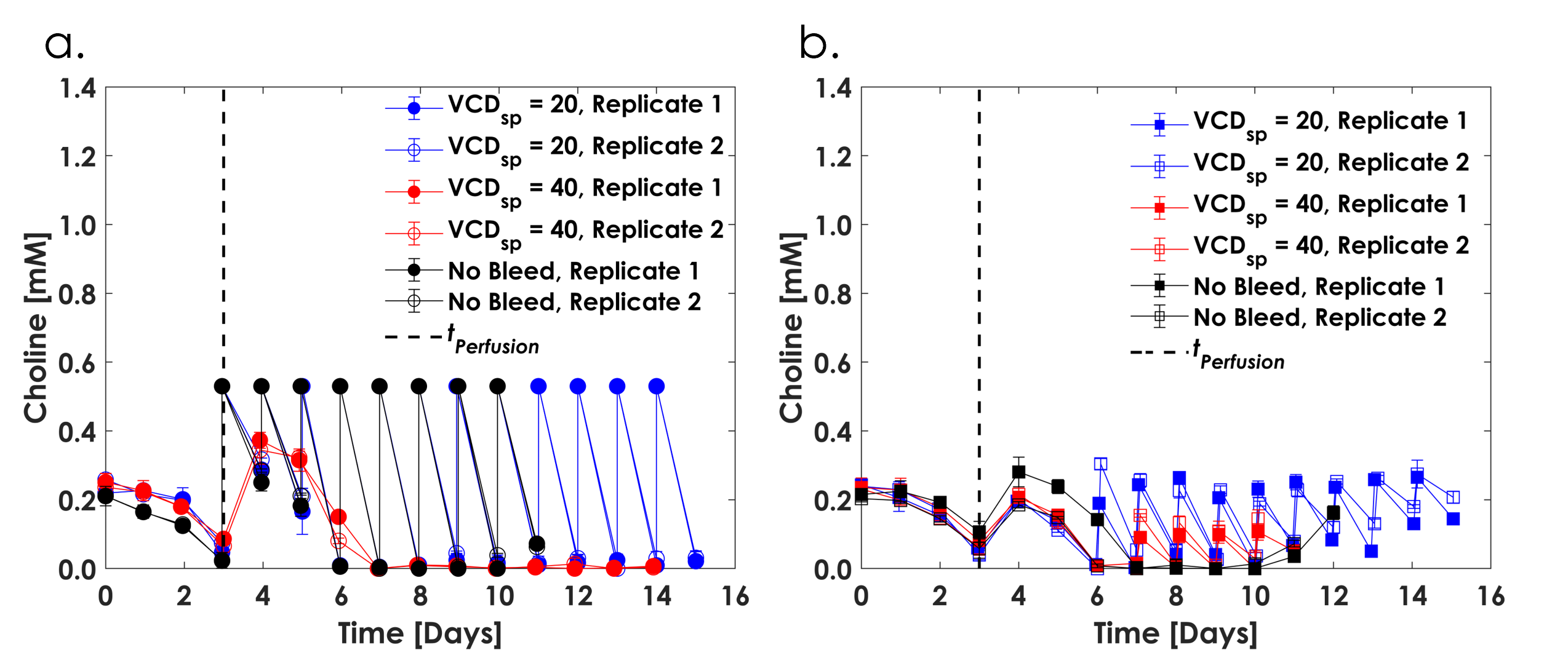

8. **Specific glycine production rates**

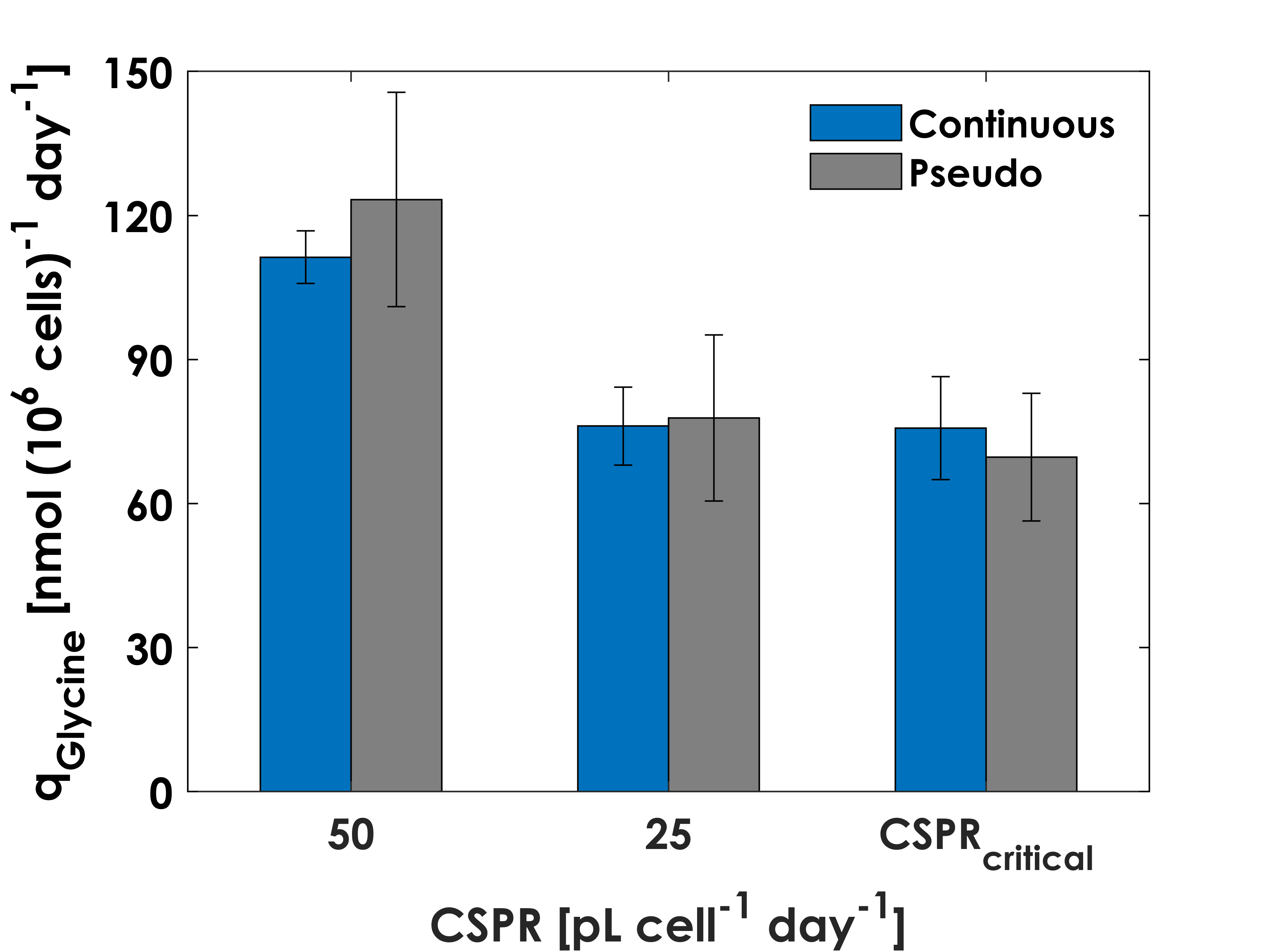

**Figure S7:** Specific glycine production rates in continuous and pseudo perfusion. Values and error bars are calculated as described in Figure 6 of the main text. n = 2 per scale and condition.

9. **The relationship between cell size, cell cycle fractions, and specific productivity**

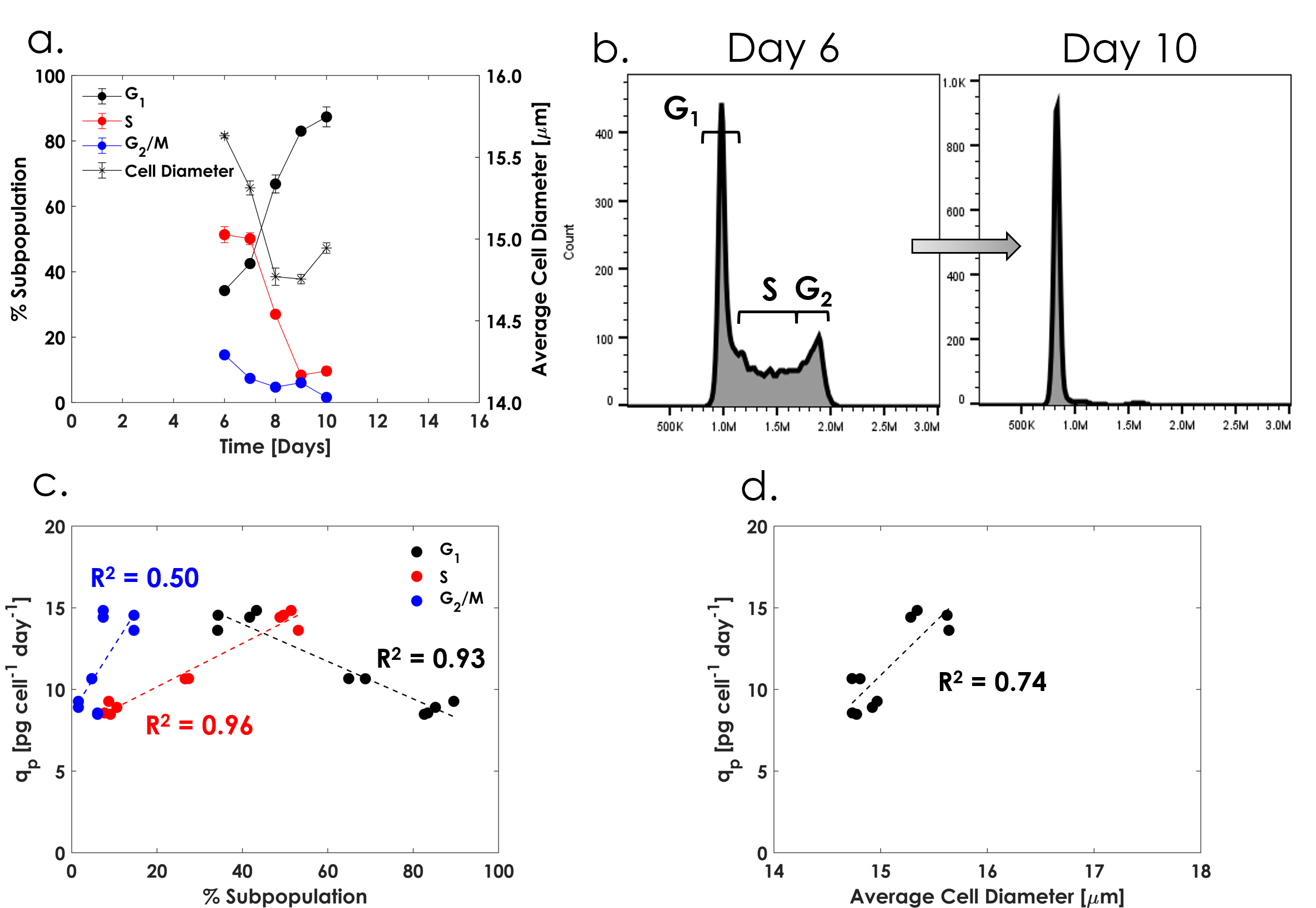

**Figure** **S8:** Pseudo perfusion with no cell bleed and perfusion rate of 1 vvd^-1^. **a)** Cell cycle fractions corresponding to the VCD profile in Figure S5.j. Values represent the mean and range of n=2. **b)** Redistribution of cell cycle fractions from day six to day ten of culture. The subpopulation gates are drawn qualitatively here for visualization but stringently determined as explained in section 2.2.2. **c)** Relationship between average cell-specific productivity and cell cycle fractions. Data from individual biological duplicates from day six to ten are shown. **d)** Relationship between cell-specific productivity and the average cell diameter. Data from individual biological duplicates from day six to ten are shown.

Fed-batch literature has consistently demonstrated that under conditions promoting a G_1_ majority subpopulation via cell cycle arrest in the G_1_ phase through temperature reductions [2], addition of small molecule cell cycle inhibitors [3], or feed supplementation with fatty acids [4], an increase in cell-specific productivity is obtained yet with varying impacts to mAb product quality profiles. Under the conditions explored in this work and for this cell line, cell-specific productivity appears growth-related given the decreasing productivity with decreasing growth rate (Figure 6.a, 6.b in the main text). Relative to the cell cycle phase distributions, increasing cell-specific productivity negatively correlates with the G_1_ phase and positively correlates with the S and G_2_/M phase subpopulations (Figure S8.d) which has also been previously reported for CHO cells [5]. Extensive work has been done to show that the increase in specific productivity in fed-batch cultures is driven by an increase in the cell size, which is also observed in this work (Figure S8.c), with the apparent correlation to any specific cell cycle phase being a secondary effect [6-8]. While [7] demonstrated a positive relationship between the cell-specific productivity and the G_1_ phase for a fed-batch process with an IgG1-producing CHO cell line, and for an unreported perfusion process, they further observed an increase in the size of the G_1_ cells themselves over the culture duration leading to the increase in specific productivity. Transcriptomic analysis suggested that the mechanisms behind the G_1_ arrest and subsequent increase in cell size were independent. These findings imply that both the distribution of cell cycle subpopulations and the average cell size are important parameters to consider in perfusion process design to reduce medium utilization while boosting productivity at the desired operating point.

The average cell diameter in this study decreases moderately with increasing VCD for a fixed perfusion rate concomitant with an increase in the G_1_ subpopulation (Figure S8.a, S8.b). As cells actively progress through the cell cycle, the average cell diameter increases with each phase and is largest during the G_2_/M phase before cell division [6], in part explaining the positive correlations observed in this work. Apart from the G_1_ majority subpopulation at lower CSPRs, which inherently reduces the average cell size, the decrease in average cell size observed here may also arise from the depletion of biomass precursors and other nutrients required for cell maintenance [7, 9], such as choline (Figure S6).
